# Microbial consortiums of putative degraders of low-density polyethylene-associated compounds in the ocean

**DOI:** 10.1101/2021.11.23.469798

**Authors:** Maria Pinto, Zihao Zhao, Katja Klun, Eugen Libowitzky, Gerhard J Herndl

## Abstract

Polyethylene (PE) is one of the most abundant plastics in the ocean. The development of a biofilm on PE in the ocean has been reported, yet whether some of the biofilm-forming organisms can biodegrade this plastic in the environment remains unknown. Via metagenomics analysis, we taxonomically and functionally analysed three biofilm communities using low-density polyethylene (LDPE) as their sole carbon source for two years. Several of the taxa that increased in relative abundance over time were closely related to known degraders of alkane and other hydrocarbons. Alkane degradation has been proposed to be involved in PE degradation, and most of the organisms increasing in relative abundance over time harboured genes encoding proteins essential in alkane degradation, such as the genes alkB and CYP153, encoding an alkane monooxygenase and a cytochrome P450 alkane hydroxylase. Weight loss of PE sheets when incubated with these communities and chemical and electron microscopic analyses provided evidence for alteration of the PE surface over time. Taken together, these results provide evidence for the utilization of LDPE-associated compounds by the prokaryotic communities. This study identifies a group of genes potentially involved in the degradation of the LDPE polymeric structure and/or associated plastic additives in the ocean and describes a phylogenetically diverse community of plastic biofilm-dwelling microbes with the potential of utilizing LDPE-associated compounds as carbon and energy source.

**Importance:** Low-density polyethylene (LDPE) is one of the most used plastics worldwide and a large portion of it ends up in the ocean. Very little is known about its fate in the ocean and whether it can be biodegraded by microorganisms. By combining 2-year incubations with metagenomics, respiration measurements and LDPE surface analysis, we identified bacteria and associated genes and metabolic pathways potentially involved in LDPE biodegradation. After two years of incubation, two of the microbial communities exhibited a very similar taxonomic composition mediating changes to the LDPE pieces they were incubated with. We provide evidence that there are plastic-biofilm dwelling bacteria in the ocean, that might have the potential to degrade LDPE-associated compounds, and that alkane degradation pathways might be involved.

## Introduction

Over the last decades plastic production and consequently plastic pollution in the ocean have drastically increased, with 4.8 to 12.7 million tons of plastic estimated to enter the ocean each year ^1^. Despite the urgent need for understanding the impact of plastics on marine ecosystems and the growing body of literature on this subject, the fate of plastics in the oceans remains largely enigmatic.

Once in the sea, plastics are rapidly colonized by a diverse microbial community ^2^. Several studies have focused on describing the taxonomic composition of these communities and the factors that might be responsible for their succession and development ^3–12^. Microorganisms related to hydrocarbon-degraders have been found to be relatively more abundant on specific plastic types than on other plastics and non-plastic surfaces ^13–15^, especially at the early stages of biofilm formation ^8^. However, the reasons provoking these differences in the colonization pattern of different plastic types remain enigmatic. The physical and chemical properties of the plastics and the presence of additives might play an important role in the initial colonization of plastics by microbes and the subsequent microbial biofilm development ^5^. However, recent evidence also suggests that plastic biofilm communities are mostly shaped by biogeographic factors ^16^. Despite these uncertainties, however, there is growing evidence that some organisms forming biofilms on plastic in the ocean do have the ability of degrading plastic-associated compounds^17, 18^.

Polyethylene (PE) is the most widely used plastic worldwide. In the year 2018, polyethylene resin demand was estimated to make up 36% of the total nonfiber plastic production ^19^. At sea, together with polypropylene, PE is usually the most abundant plastic at the sea surface ^1, 20^.

PE is considered to be highly resistant to biodegradation, especially in the absence of previous weathering by, for instance, photo-degradation. However, studies have reported weight loss and changes in the physical and chemical structure of PE when incubated with specific microorganisms. For instance, bacteria of the genera *Bacillus* and *Pseudomonas* have been associated with PE degradation both in the marine environment ^21–24^ and in soil ^25–27^. Some studies have also suggested the implication of the alkB gene and alkane degradation pathways in PE biodegradation ^27, 28^.

This alkB gene encodes an alkane-monooxygenase that oxidizes the initial terminal end of n-alkanes. Because the main molecular structure of PE polymers is similar to alkanes, it has been suggested that the metabolism responsible of PE bio-degradation is similar to that of hydrocarbons, specifically to alkanes ^29^. Alkanes are degraded through a succession of oxidation steps leading to beta-oxidation and consequently, to the entering of acetyl-CoA into the citric acid cycle. The first oxidation step is the initial terminal hydroxylation of the alkane chains, which can be carried out by enzymes belonging to different families, usually depending on the alkane chain length ^30^. In organisms degrading short-chained alkanes, it is usually carried out by methane monooxygenase. The initial hydroxylation of medium-chained alkanes (C5-C17) is mediated by organisms that frequently contain soluble cytochromes P450, typically the cytochrome P450 CYP153 alkane hydroxylase, and integral membrane non-heme iron monooxygenases, typically alkB. *Alcanivorax borkumensis*, for example, has genes encoding for both types of medium-chained alkane monooxygenases in its genome ^31^. Organisms capable of hydroxylating long-chained alkanes (>C18), such as *Alcanivorax dieselolei* and *Geobacillus thermoleovorans*, have been shown to have the hydroxylases AlmA and LadA, respectively ^32, 33^. Except for the alkB encoded monooxygenase, any other potential enzyme or pathway that might be involved in PE biodegradation is only poorly studied, particularly in the marine environment.

In this study we investigated the potential of three PE biofilm communities (hereafter called communities A, B and C), initially attached to three different plastics collected from the Northern Adriatic Sea, to metabolize LDPE and/or associated compounds. The term LDPE-associated compounds used hereafter, encompasses the LDPE polymeric chain and any potential additives associated with the used LDPE pieces. The communities were incubated in artificial seawater (ASW) amended with inorganic nutrients and with LDPE as their sole carbon source for two years. To identify which taxa and metabolic pathways became enriched during the incubation, metagenomes were analyzed after one and two years. Oxygen consumption measurements were performed as a metabolic activity parameter to determine whether the microbial communities incubated with LDPE are more active than communities on inert surfaces such as glass. Weighing of the plastic, scanning electron microscopy (SEM) and attenuated total reflection Fourier-transform infrared (ATR-FTIR) spectroscopy were performed on different types of PE sheets to determine whether any physical and/or chemical changes to the surface of the plastic occurred due to the presence of the microbial communities (see Fig. 1 for experimental setup).

**Fig. 1.**
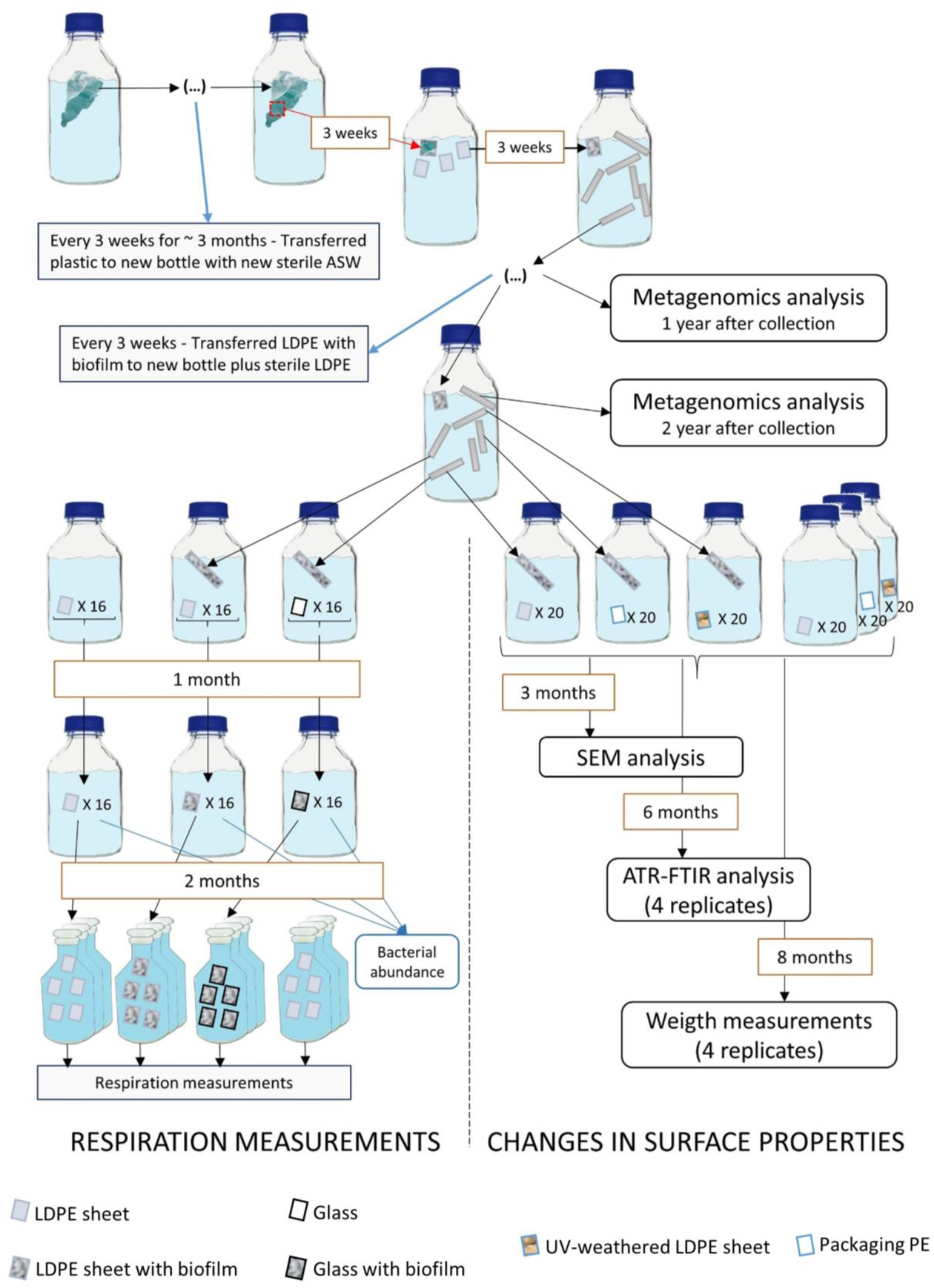
Scheme of the experimental setup. All incubations were performed in sterile artificial seawater amended with 2 µM of NaPO_4_ and 10 µM of NH_4_Cl. All bottles were kept in the dark at room temperature (∼21°C). This workflow was performed for each of the three plastics, A, B and C.

## Results

### Microbial community composition

In total, we obtained six metagenomes, one from the original community A (community A0), one for each of the communities after one year of incubation (communities A1, B1 and C1) and one for each of the communities A and C after two years of incubation (communities A2 and C2, respectively). mOTU (operational taxonomic unit obtained with the mOTUs software) diversity and richness of communities A and C decreased from the original community (A0) to after one year of incubation in the case of community A (A1), and after two years of incubation in the communities A and C (A2 and C2) (Fig. S1).

Analysis of the 16S rRNA gene assembled from the metagenomes revealed that after two years of incubation both, community A2 and C2 exhibited a similar taxonomic composition at the family level (Fig. S2) with *Phycisphaeraceae*, *Planctomycetaceae*, *Rhodobacteraceae*, *Rhodospirillaceae* and *Saprospiraceae* representing similar relative abundances and constituting a major fraction of the classified families (Fig. S2). The *Rhodobacteraceae* was the most abundant of all the identified families in communities A2 and C2 with relative abundances of 16% and 20%, respectively. *Rhodobacteraceae* were also abundant in community B1 with a relative abundance of 7%. It is nevertheless important to note that 55%, 64%, 68%, 73% and 70% of the families classified with matam from communities A1, A2, B1, C1, C2, respectively, remained unclassified.

### Phylogenetic classification of metagenome assembled genomes (MAGs)

From the 733,317 co-assembled contigs, a total of 61 MAGs with >50% completeness and <10% contamination was recovered (Table S1). The % of bps from the total reads of each sample assigned to the identified MAGs varied greatly between samples with 3.3%, 58.9%, 78.5%, 24.6%, 73.2% and 81.9% of the total bps of the metagenomes from communities A0, A1, A2, B1, C1 and C2, respectively, assigned to the bps of all 61 MAGs. Sixty of the MAGs were classified as bacterial and one as archaeal (MAG 53) (Fig. 2). The archaeal bin was classified as *Crenarchaeota* and was most abundant in community C1, but was not found anymore in C2, and was also present in very low relative abundances in communities B1 and A2 (Fig. 2, Table S1).

**Fig. 2.**
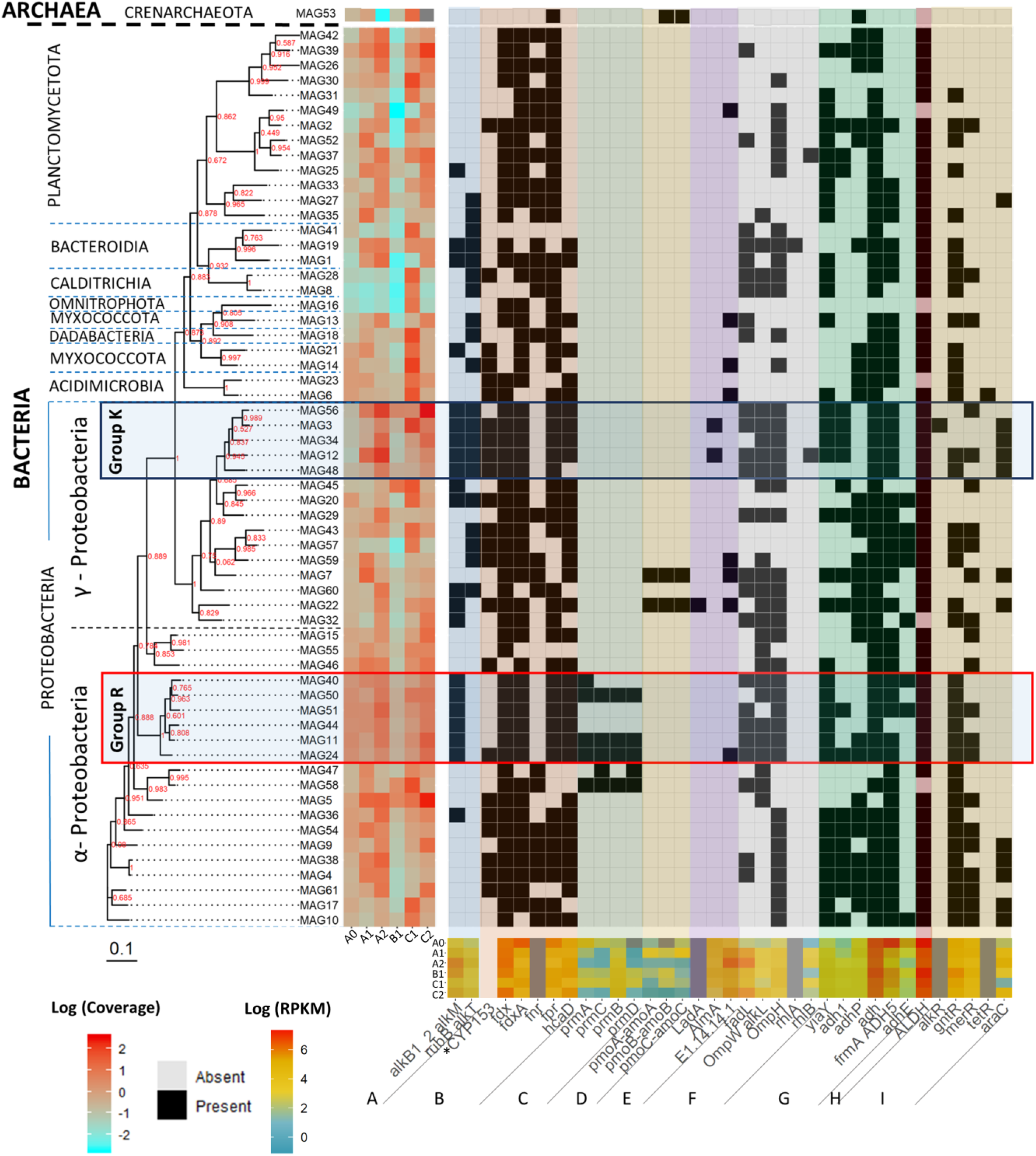
Phylogenetic tree of the identified MAGs, their coverage in the six metagenomes plus the presence or absence of selected alkane and/or fatty acid degradation related genes (Table S2) in each of the MAGs and their RPKM (reads per kilobase million) in the different samples. The tree was built with Fasttree using the multiple sequence alignment obtained with gtdb-tk. Bootstrap values of each tree branch are given in red. The different gene groups refer to: (A) genes related to the alkane 1-monooxygenase-encoding gene alkB; (B) genes related to the cytochrome P450 alkane hydroxylase CYP153; (C) a propane oxygenase encoding genes; (D) a methane monooxygenase encoding genes; (E) genes encoding long-chain alkane related monooxygenases and fatty acid oxygenases; (F) transporters and biosurfactant synthesis genes; (G) alcohol dehydrogenase genes; (H) aldehyde dehydrogenase genes; and (I) genes encoding for proteins thought to be involved in transcription regulation of alkane degradation. Group K is a taxonomic clade, classified as *Pseudomonadales* that includes the MAGs that had a high number of the selected genes and were closely related to *Ketobacter* sp. and *Alcanivorax* sp. Group R includes all MAGs classified as *Rhodobacteraceae*, which all have the alkB gene encoding the alkane monooxygenase most typically associated with alkane degradation. KEGG annotations were used to determine RPKM values of genes in different samples. *The CYP153 gene does not have an assigned KO identifier, its presence/absence in MAGs was identified using hmm models.

MAGs 5 and 56, belonging to the family *Parvibaculaceae* and genus *Ketobacter*, respectively, increased in coverage in both, community A and C from the first to the second year, and were among the MAGs with the highest coverage in the communities A2, B1 and C2 (Fig. 2, Table S1). We compared the obtained MAGs with the genomes of potential hydrocarbon, alkane or plastic degraders (Table S3) retrieved from NCBI (Fig. 3A). MAGs 5 and 56 clustered together with alkane degrading species of the genus *Parvibaculum* and alkane degrader *Ketobacter alkanivorans*, respectively (Fig. 3A). MAG 56 was grouped in a larger cluster (Group K in Figs. 2 and 3) composed of MAGs classified as *Alcanivoraceae* and *Ketobacteraceae* and the genomes of known *Alvanivorax* species and *Ketobacter alkanivorans* (Fig. 3). Generally, MAGs from Group K increased over time (Fig. 2). Other MAGs also increased in coverage over time in the communities A and C (MAGs 1, 13, 15, 26, 27, 37, 39, 42, 44, 49, 51, 60, 61) (Fig. 2). However, a differential expression analysis determined that only MAGs 9, 11, 15, 24, 29, 32, 34, 56, 60 and 61 had genes that were significantly enriched after two years of incubation in comparison to the one-year incubations in communities A and C (Table S4).

**Fig. 3.**
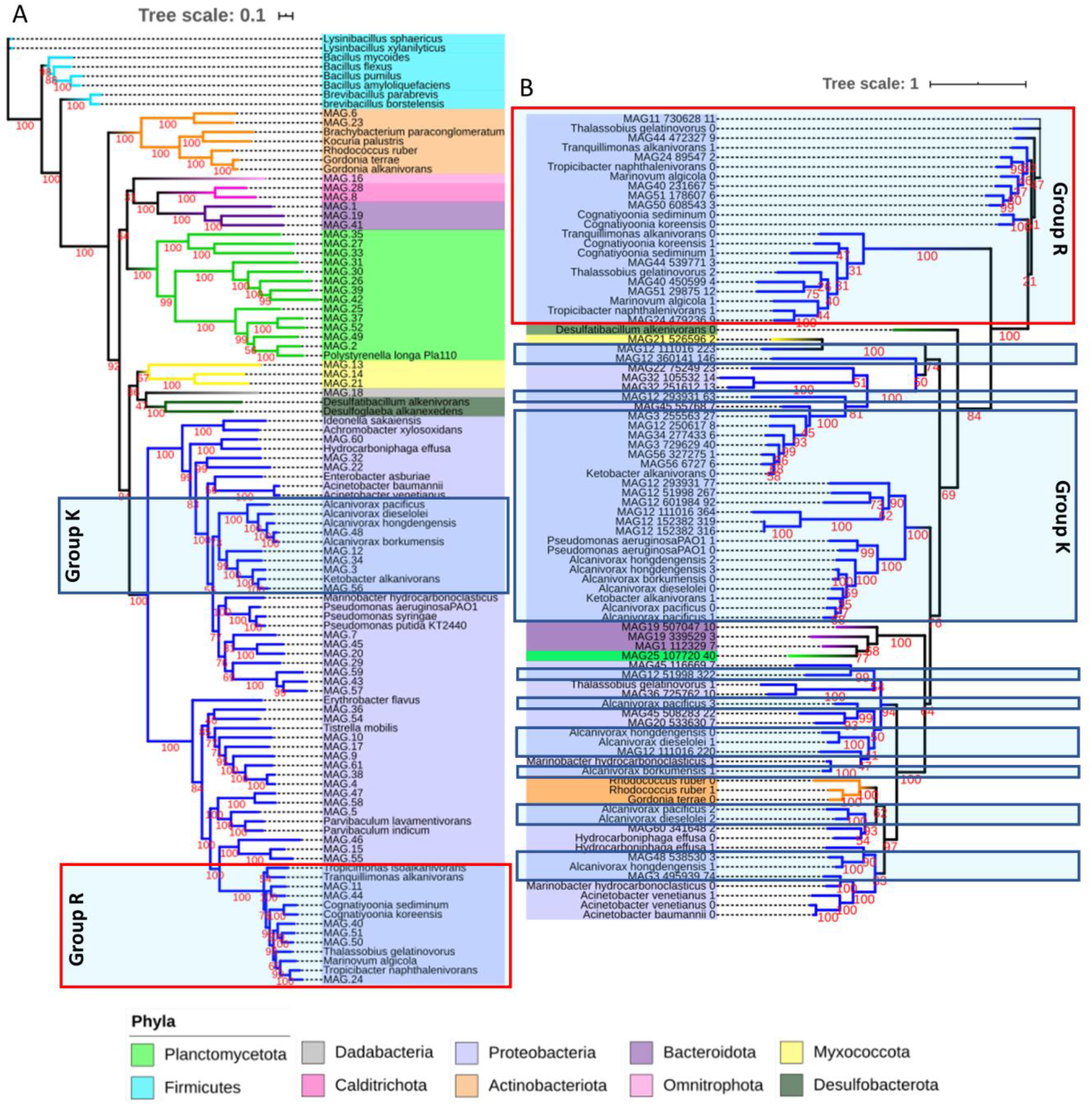
Phylogenetic placement of (A) the MAGs and selected genomes (Table S3) and (B) putative alkB sequences identified in MAGs and genomes. Bootstrap values are indicated in red. Group K (blue boxes) and Group R (red boxes) refer to the same groups as described in Fig. 2.

Most MAGs exhibited a very low coverage in community B1, with the exception of some classified as *Proteobacteria*, which is also supported by the 16S rRNA gene analysis (Fig. S2). MAGs 5, 58, 45 and 56 were the only ones with coverage values higher than 1 in community B1, with values of 3.93, 2.95, 12.57 and 5.11, respectively.

### Functional gene dominance in the metagenome over time

A total of 460 genes were significantly enriched (Table S4) after two years of incubation in comparison to the one-year incubations in communities A and C as determined by a differential expression analysis. Ten, 108, 4, 17, 10, 2, 204, 1, 21 and 2 of these genes were respectively assigned to MAGs 9, 11, 15, 24, 29, 32, 34, 56, 60 and 61. The remaining 81 genes were not assigned to any MAG.

A Deseq2 analysis of all genes when grouped in KEGG orthologs (KO numbers) showed a significant decrease in relative abundance (p-value < 0.05) of certain selected genes (Table S2), namely genes araC, pmoA-amoA and genes encoding for a Propane 2-monooxygenase (prmA, prmC and prmD) from one year of incubation to two years of incubation in communities A and C.

To determine which metabolic pathways increased in the incubations over time, we compared the predicted genes with RPKM (reads per kilobase million) values that increased from the original community to after one year of incubation in the case of community A, and from one to two years of incubation in both, communities A and C. Predicted genes annotated as GST (glutathione S-transferases) homologues were overall the most abundant genes increasing in RPKM over time (Table S3), even though they were already present in fairly high relative abundance in the original sample A0 (Table S4). GST homologue genes are involved in PAH (polycyclic aromatic hydrocarbons) degradation in prokaryotes, as well as in the biodegradation of other xenobiotics ^34^. Other enriched genes in the LDPE treatments, like atzF and mlhB, are also involved in xenobiotics degradation.

A large part of the oxidoreductase-encoding genes increasing in RPKM in the communities A2 and C2 were annotated as genes related to lipid metabolism, especially fatty acid and steroid metabolism (Table S5). Furthermore, all the identified complete KEGG pathway modules with increasing RPKM values were related to lipid metabolism, particularly fatty acid biosynthesis, fatty acid beta-oxidation with the synthesis of acyl-CoA, and acylglycerol and sphingosine degradation. The CYP51 gene (annotated by KEGG) was the gene of the oxidoreductases category that increased most pronouncedly in RPKM in community A2 and C2 over time (Fig. S3). It encodes sterol 14 alpha-demethylase, a cytochrome P450 enzyme which plays an important role in sterol synthesis and also in fatty acid metabolism ^35^. Another oxidoreductase, cypD_E, also significantly increased in RPKM over time, especially in community C2 (Fig. S3). CypD_E encodes for cytochrome P450/NADPH-cytochrome P450 reductase linked to fatty acid oxidation. It catalyzes the hydroxylation of a range of long-chain fatty acids, oxidises reduced nicotinamide adenine dinucleotide phosphate (NADPH) by electron transfer to the heme iron of the cytochrome P450 N-terminal domain ^36^.

Two predicted genes, annotated as monooxygenases involved in fatty acid degradation, also increased in RPKM over time in the communities A2 and C2. While one of the predicted genes was classified encoding an unspecific monooxygenase (EC:1.14.14.1, KO identification K00493), the other gene was annotated as gene alkB1_2, encoding an alkane 1-monooxygenase involved in alkane degradation (Fig. 1 and Fig. S3). Alkane 1-monooxygenases catalyse the hydroxylation of n-alkanes and fatty acids in the presence of NADH-rubredoxin reductase and rubredoxin. A gene annotated as rub, encoding the rubredoxin-NAD(+) reductase, was also among the most enriched genes over time (Fig. 2, Tables S5 & S6). The rubredoxin-NAD(+) reductase transfers electrons from NADH to rubredoxin reductase and further through rubredoxin to alkane 1-monooxygenase.

There were several other genes increasing over time encoding enzymes related to fatty acid metabolism. Some of these genes, i.e., genes encoding lipid and fatty acid transporters (genes lip, fadL and OmpW) and alcohol (genes yiaY, adh1) and aldehyde (gene ALDH7A1) dehydrogenases (Tables S5 & S6)^30^ are the key genes in the alkane degradation pathway.

### Putative hydrocarbon degradation genes in MAGs

Overall, most MAGs that increased in coverage over time either showed the potential to degrade alkanes or had genes encoding for alkane and hydrocarbon transporters. MAGs 3, 12 and 22 had at least one medium-length alkane oxidation related gene and either the AlmA or the LadA gene, encoding enzymes potentially degrading both long- and medium-length chain alkanes (Fig. 2). MAG 22 also had all three genes encoding a methane/ammonia monooxygenase, suggesting that this organism is also capable of degrading short-chain alkanes. Furthermore, these MAGs also had at least one gene encoding a surfactant and alkane membrane transporters. Group K MAGs (shown in Fig. 2) all had genes encoding aldehyde and alcohol dehydrogenases, at least two alkane related transporters and proteins associated with surfactant production. They also harboured the alkB1_2 and alkT gene encoding alkane monooxygenase and rubredoxin---NAD+ reductase, respectively, which together form a complex capable of oxidizing medium-chained alkanes (Fig. 2). Group K MAGs also harboured genes encoding a cytochrome P450 alkane monooxygenase (CYP153), the associated transcription regulator gene araC and the genes encoding a ferredoxin and ferredoxin reductase, which are required for the transfer of electrons from NAD(P)H to the cytochrome ^37^.

MAG 5, which is amongst the most abundant MAGs in community C2, harboured a gene encoding a protein associated with surfactant production and alkane transport into the cell (OmpW, alkL). MAGs 24, 11, 44, 51, 50 and 40, which together formed the *Rhodobactereaceae* clade (Group R in Fig. 2) all had genes annotated as alkB1_2, hcaD and the surfactant producing membrane protein-encoding gene OmpH. No rubredoxin reductase encoding gene was annotated in the Group R MAGs. However, they all had genes encoding for ferredoxin and ferredoxin reductase. It has been suggested that ferredoxin and ferredoxin reductase might replace rubredoxin and rubredoxin reductase usually associated with alkB ^38^. Together with MAG 58, three of the Group R MAGs also harboured the four genes encoding for the different subunits (Fig. 2, prmABCD) of an oxidoreductase that oxidises propane (propane 2-monooxygenase), a three-carbon alkane that is a gas at standard temperature and pressure.

Furthermore, group R MAGs, except MAG 51, harboured the gene OmpW/alkL encoding an outer membrane protein, involved in 1-alkanes transport into the cell and surfactant production. MAG 24 also had a gene annotated as an unspecific monooxygenase (E1.14.14.1). With the exception of MAG 50, all other Group R MAGs harboured the gene fadL. Almost all MAGs harboured at least one aldehyde dehydrogenase and one transcriptional regulator encoding gene, both involved in several metabolic processes within the cell.

### Distribution of the alkB and CYP153 gene in the MAGs

HMM models were used to identify putative alkB and CYP153 genes in the MAGs and the genomes of potential hydrocarbon and plastic degraders (Table S3). A total of 21 and 23 MAGs had at least one copy of genes classified as alkB and CYP153, respectively (Fig. 2). AlkB was found in one *Myxococcota* and one *Planctomycetota* MAG, in two *Bacteroidota* and in seventeen *Proteobacteria* MAGs. Multiple copies of the alkB gene were found in some of the MAGs, especially in MAGs from Group K. The highest number of alkB genes in one MAG was 12, found in MAG 12 (Fig. 3B). AlkB genes belonging to Group R MAGs clustered together in one single group. The alkB genes from Group R MAGs clustered with the alkB genes from known marine *Rhodobacteraceae* alkane degraders (Fig. 3B). In contrast, alkB genes from Group K clustered together in several groups, with some of them grouping with sequences from other *Proteobacteria* MAGs (Fig. 3B).

CYP153 was detected in one MAG of *Calditrichota* and of *Plactomycetota*, in two MAGS of *Actinobacteriota* and of *Myxococcota* and in seventeen MAGs classified as *Proteobacteria*. There were several clusters composed only of *Proteobacteria* CYP153, however, there were also CYP153 genes from MAGs of different phyla clustering together (Fig. S4). There were several MAGs with more than one copy of a gene classified as CYP153 (Fig. S4).

### Cell abundance and respiration measurements

To determine respiration rates of the biofilms on plastic in comparison to glass, oxygen consumption measurements were performed after the surfaces were incubated for three months with communities A2, B2 and C2 (Fig. 1). Just before starting the respiration measurements, the abundance of all three microbial communities was significantly higher in the incubations with plastics than with glass (F=955.18, p<0.001) (Fig. 4B and Fig. S5). The cell abundance of the three communities growing on LDPE also differed between each other (F=96.41, p<0.001, Table S6), with community C exhibiting a significantly higher number of cells than the communities A and B (Fig. 4). No cells were found on the surface of both LDPE controls (Control I and Control II).

**Fig. 4.**
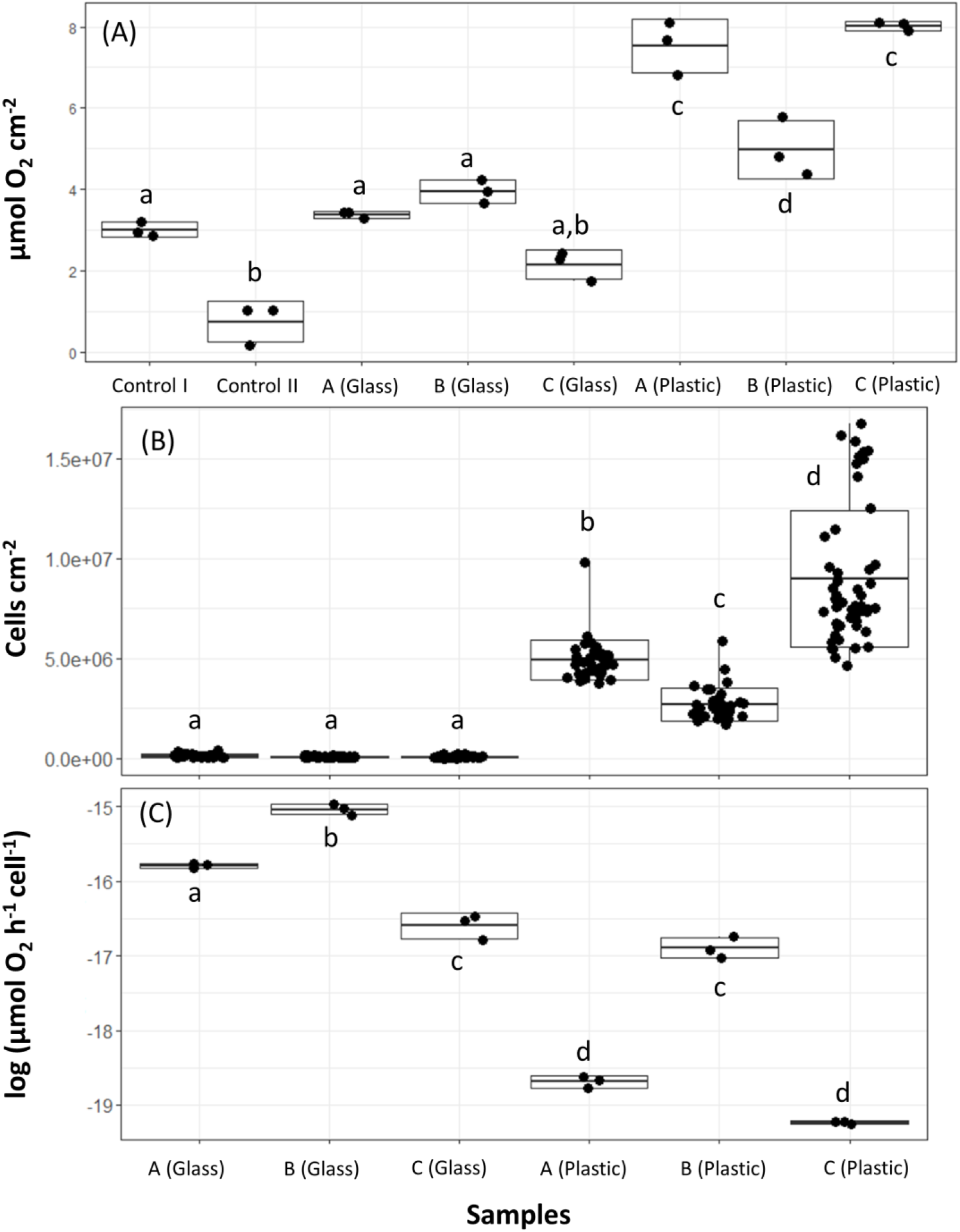
Oxygen consumption rates of microbes in the plastic and glass treatments. The boxplots represent the mean ± SD. (A) Total oxygen consumption in each sample in µmol per cm^2^ of plastic or glass. Each black dot represents one replicate. (B) Number of cells per cm^2^ of plastic or glass. The boxplot lines in A, B and C (Plastic) represent the maximum number of cells cm^-2^ counted in a photo of that sample. Each black dot corresponds to the number of cells per cm^2^ of one photo. The controls were excluded from this and the next plot because no cells in those samples were detected. (C) Oxygen consumption cell^-1^ h^-1^. Samples with different letters on each plot were significantly different from each other (p-value < 0.05, Tukey test results in Tables S7-9).

Oxygen consumption rates per cm^2^ surface area were significantly higher in the microbial communities associated with LDPE than on glass (F=48.84, p<0.001) and in the controls (Fig. 4A, Table S7). However, cell-specific oxygen consumption rates were higher in microbial communities in the glass than in the plastic treatment (Fig. 4C). Incubations with community B2 showed the least difference in bulk oxygen consumption rates between the glass and plastic treatment. Additionally, from all the treatments with LDPE, community B2 exhibited the highest cell-specific oxygen consumption rate. The communities A2 and C2 showed similar cell-specific oxygen consumption rates (Fig. 4C, p=0.99). However, when incubated with glass, community A2 showed a higher oxygen consumption rate than community C2 (Fig. 4C, p<0.001).

### Alteration of plastic surfaces determined by ATR-FTIR, SEM and weight

The ATR-FTIR spectra of the untreated LDPE sheets, packaging LDPE and UV-treated LDPE sheets (prior to microbial incubations) and after six months of exposure to microbial communities growing previously on PE as sole carbon and energy source for two years are depicted in Fig. 5. All plastics shared the typical LDPE bands for CH_3_ and –CH_2_– groups at 2916, 2849, 1469, 1375 and 718 cm^-1^.

**Fig. 5.**
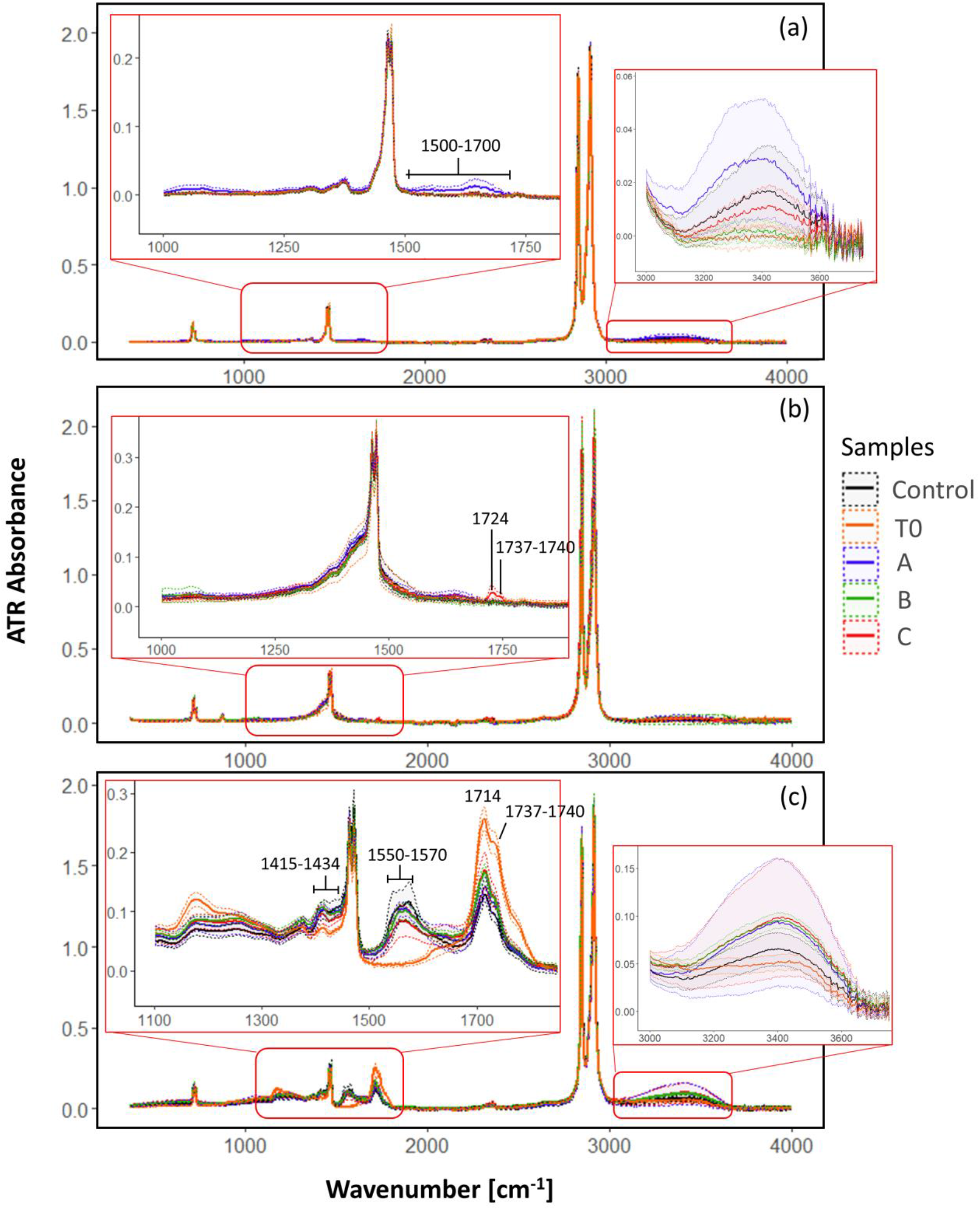
ATR-FTIR spectra of (a) non-weathered LDPE sheets; (b) packaging PE sheets; and (c) UV-weathered LDPE sheets prior to the incubation and after six months incubated with the three microbial communities. The lines correspond to the mean value of the replicates and the same-coloured dashed lines represent the corresponding standard deviation.

In non-weathered LDPE plastics incubated with community A, a band spreading from 1500 to 1700 cm^-1^ appeared indicating the presence of carbonyl (>C=O) and vinyl groups (– CH=CH–). Also, a small peak at 3400 cm^-1^ was detected which might indicate the appearance of hydroxyl groups. Bands at around 1640 and 3400 cm^-1^, however, might also indicate the presence of water. The spectra from packaging PE pieces incubated with community C revealed peaks at 1724 and at 1737-1740 cm^-1^, corresponding to the appearance of carbonyl groups, carboxylic groups and aldehydes, respectively. At the beginning of the incubations, UV-weathered plastics exhibited an additional band at 1714 cm^-1^, corresponding to the appearance of carbonyl groups resulting from photo-oxidation of polyethylene. After six months of incubation, peaks indicating carboxylates appeared at 1550-1570 cm^-1^ and 1415-1434 cm^-1^ in all incubations. The peak at 1714 cm^-1^ had higher intensity in UV-weathered plastics incubated with microbes than in the control samples without microbes, but was of lower intensity than prior to the incubation experiment. A small peak at about 3400 cm^-1^ also appeared, especially in the plastics incubated with bacteria. There were also signs of degradation in the control treatment without microbes of the UV-weathered LDPE. This degradation might be the result of the presence of inorganic ions working as oxidizing catalysts of the carbonyl compounds to carboxylic compounds ^39^.

After three months of incubation a visible biofilm developed on all PE samples, except in the controls (Fig. 6A-F). In some samples, after removal of the biofilm, there were some visible holes and apparent changes in the plastic surface, especially in the previously UV-weathered LDPE and packaging PE (Fig. 6G-I) when incubated with the community A and C.

**Fig. 6.**
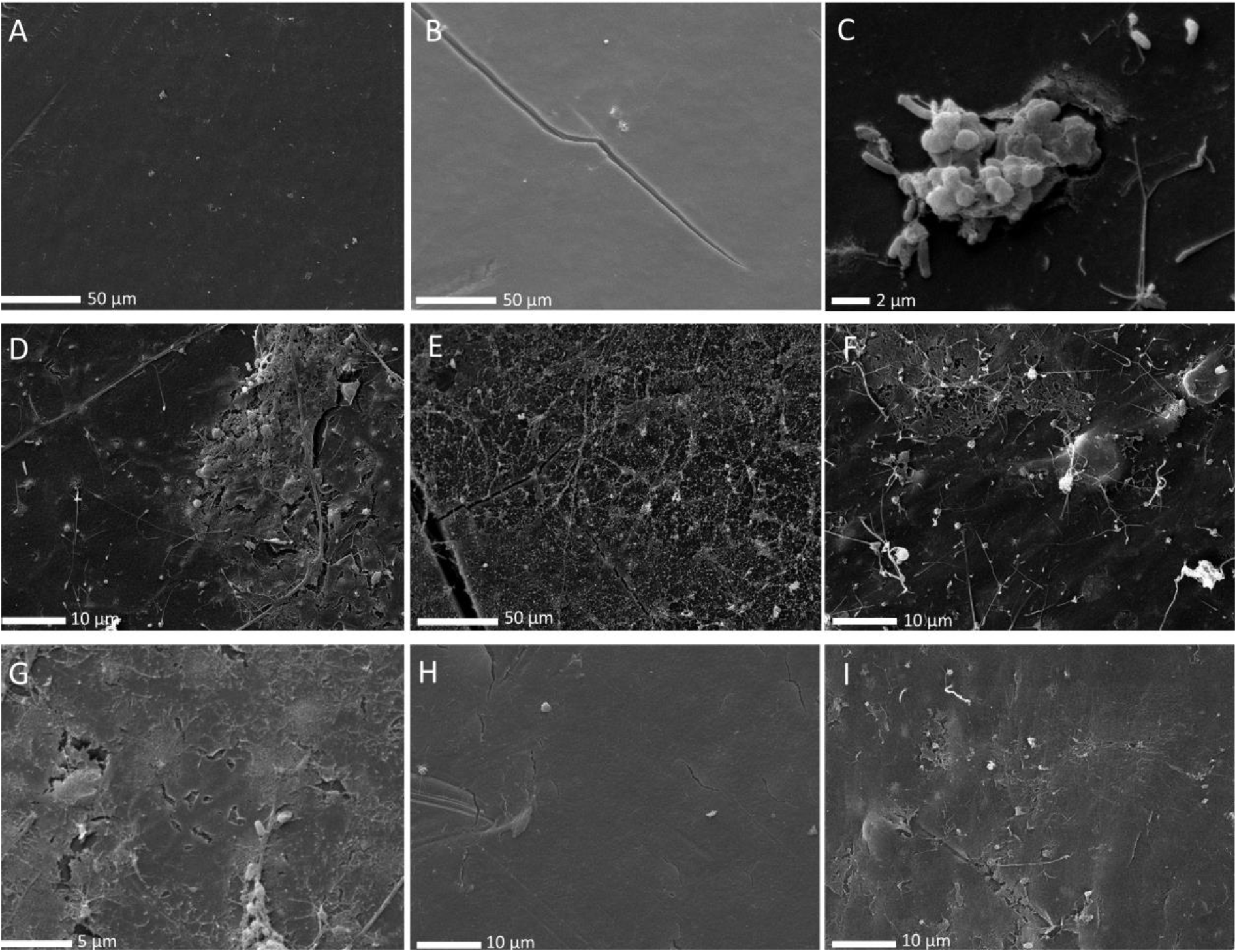
SEM images of samples collected after three months of incubation with the three different communities. (A) LDPE control sheet; (B) UV-weathered LDPE control; (C) LDPE sheet incubated with community C2; (D) LDPE sheet incubated with community A2; (E) UV-weathered LDPE incubated with community C2; (F) Packaging PE incubated with community C2; (G) LDPE sheet incubated with C2, after biofilm removal; (H) UV-weathered LDPE sheet incubated with B1 after biofilm removal; (I) Packaging PE incubated with C2 after biofilm removal. The white bars represent the scale of the image.

Significant weight losses were found between different treatments and/or time points in the packaging PE and non-weathered LDPE (F=6.847, p<0.001 and F=9.34, p<0.001 respectively). Non-weathered LDPE plastics significantly decreased in weight over a period of eight months of incubation compared to the controls without bacteria, (Fig. 7, Table S8). The loss in weight over the eight months amounted to 7.5%, 5% and 8.5% in plastics incubated with communities A, B and C, respectively. All PE packaging plastics, including the control samples, significantly decreased in weight after eight months of incubation (Fig. 7).

**Fig. 7.**
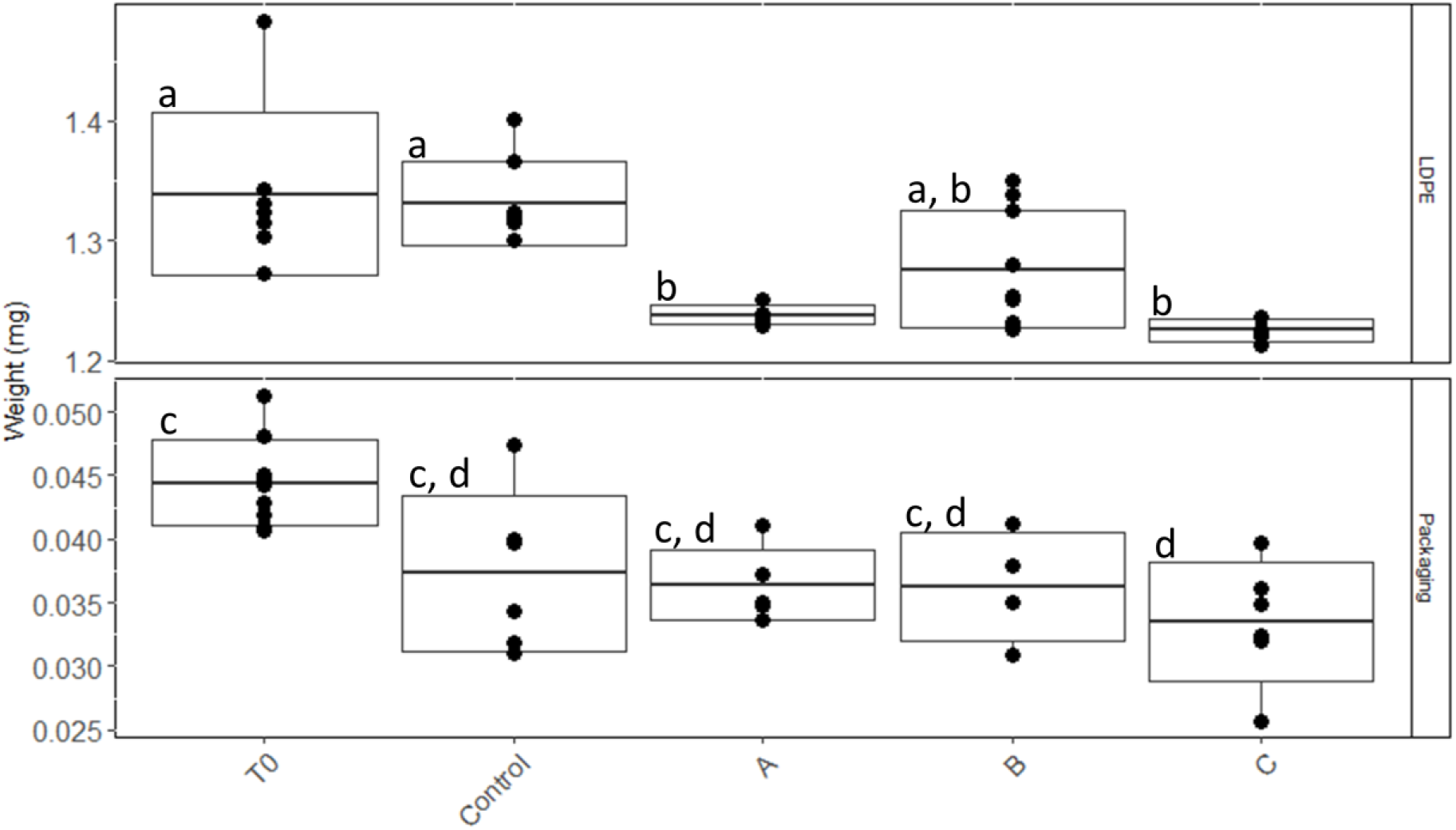
Weight of non-weathered LDPE and packaging PE pieces incubated with the three different communities A, B and C for eight months. The boxplot represents the mean ± SD with the whiskers representing the minimum and maximum weight values. Each dot represents the weight of an individual plastic piece. Different letters within each panel refer to samples with statistically significant differences (p-value < 0.05, Table S8).

## Discussion

Our results suggest that LDPE-associated compounds can serve as a carbon and energy source for at least a part of the plastic biofilm community. We have also identified several organisms, genes, and associated pathways related to alkane and fatty-acid metabolism, that might be involved in the utilization of LDPE and its associated compounds as a carbon source.

Even though the three communities were derived from three different plastics and even a different season in the case of community C, there were several similarities in taxonomy and functional diversity between the communities A and C after two years of incubation. The original community A0 was dominated by families typically found in plastic biofilms, such as *Flavobacteriaceae, Pseudoalteromonaceae*, *Alteromonadaceae* and *Rhodobacteraceae* (Fig. S2) ^40^. Over time, the families *Flavobacteriaceae, Pseudoalteromonaceae*, *Alteromonadaceae* decreased in relative abundance, while the family *Rhodobacteraceae* dominated the classified families in the communities A2 and C2 (Fig. S2).

The two MAGs with the highest coverage in the A2 and C2 communities, classified as *Parvibaculaceae* and *Ketobacter* sp. (MAGs 5 and 56, respectively), are closely related to species known to be involved in alkane degradation (Fig. 3A). *Ketobacter alkanivorans* is a recently characterized marine n-alkane degrading bacterium and, until now, the only classified species of the *Ketobacter* genus ^41^. Even though in the recently proposed rank-normalised GTDB taxonomy ^42^ *Ketobacter* belongs to the *Ketobacteraceae* family, it was previously grouped together with the genus *Alcanivorax* in the *Alcanivoraceae* family. *Alcanivorax* and closely-related bacteria have been shown to thrive after oil spills ^43^ and dominate petroleum-contaminated waters when adequate amounts of nitrogen and phosphorus are available ^44^. This is mainly attributed to their high ability in degrading alkanes. In our incubations, *Ketobacteraceae* and *Alcanivorax* were among the MAGs with the highest increase in relative abundance over time and they have the genetic potential for alkane degradation. While the role of alkane degradation in LDPE degradation is unclear, our results further suggest that if the LPDE polymer can in fact be degraded, alkane-related metabolism might be involved.

The family *Parvibaculaceae* (previously *Rhodobiaceae*) to which MAG 5 belongs to, comprises species from the genus *Parvibaculum* known for their ability to degrade polycyclic aromatic hydrocarbons and laundry detergents ^45–47^. Additionally, some of the other relatively abundant MAGs in the communities A and C after one and two years of incubation, like the ones classified as *Acidimicrobia* (previously *Actinobacteria*) and *Myxococcota* (previously order *Myxococalles*), have also been found in hydrocarbon polluted areas and associated with alkane degradation ^48^. These MAGs also harboured essential genes encoding enzymes for alkane degradation. The fact that several of the most abundant MAGs in communities A2, B1 and C2 were identified as close relatives to hydrocarbon degraders and harboured genes encoding all essential enzymes for alkane degradation is additional evidence that hydrocarbon degrading organisms were positively selected in our incubations. This supports the hypothesis that alkane degradation pathways might be used in the degradation of LDPE-associated compounds. While *Actinobacteriota* and several alkane degrading *Proteobacteria*, like those belonging to the *Alcanivorax* genus or the *Rhodobacteraceae* family, have already been identified and characterized ^49–53^, alkane degraders belonging to the *Myxococcota* and *Planctomycetota* phyla are poorly studied. Our phylogenetic analysis coupled with the identification of alkane oxygenase genes in taxonomically distant MAGs indicates that there might be a relatively high diversity of unidentified alkane degraders in the biofilm of PE in the sea. The few studies that looked into the genetic potential for PE degradation in the ocean mainly focused on the alkB gene ^28^. After two years of incubation, the high relative abundance of MAGs with other non-alkB alkane monooxygenases indicates that pathways other than those associated to alkB, might also be relevant for the utilization of PE-originating compounds and should be considered in future research.

Even after two years of incubation, some of the MAGs do apparently not have the genes required for alkane oxidation or transport into the cell. This suggests that there might be other non-identified pathways involved in the utilization of LDPE derived carbon. This could, for example, explain the increase in relative abundance of several *Planctomycetota* MAGs in which no alkane oxidase was found. While two of the *Planctomycetota* MAGs had genes classified as alkane oxygenases, the majority did not. Albeit in low relative abundances, members of this phylum have been previously found in oil-degrading consortia ^54^ and in fact, some have been reported to have alkane monooxygenase genes ^55^. As common elements of plastic biofilms ^8, 56^, the potential for members of this phylum to utilize PE originating compounds should be considered.

The lack of genes in some MAGs related to alkane degradation, however, can also be a result of the low completeness of some of these genomes, not allowing the detection of the specific genes. It might also simply reflect the fact that some of these MAGs do indeed not have the ability to degrade hydrocarbons derived from LDPE and are sustained by by-products of the hydrocarbon-degrading component of the community or other compounds potentially leaching from the plastic or generated by the biofilm associated microbes. Community B1 appears to also have been dominated by bacteria from Groups R and K and MAG 5 (Fig. 2), giving the community the necessary repertoire of genes for surfactant production and alkane degradation. However, this community showed slower biofilm development than the other two communities despite exhibiting similar diversity (Fig. S1, Fig. 4). Furthermore, non-weathered LDPE sheets lost less weight when incubated with community B than with any of the other two communities (Fig. 7). This might be a result of differences in LDPE degradation efficiency between the different organisms in each of the incubations but might also suggest that community B might have been limited in specific compounds that communities A and C were able to produce. This implies that, if the degradation of LDPE-related hydrocarbons is in fact taking place, the degraders while capable of sustaining growth of a diverse bacterial community, might be relatively slow growing and require metabolic diversity to efficiently degrade LDPE-associated compounds. This is in accordance with laboratory experiments on the degradation of untreated PE by bacterial monocultures, exhibiting low biodegradation rates ^21, 57^. Generally, the FTIR spectra revealed that changes in the surface of the PE incubated with the three communities were relatively small after six months. However, bacterially mediated modifications of the surface of the plastics, especially on non-weathered LDPE by community A and on packaging PE when incubated with community C, caused an apparent weight loss of these plastics after eight months of incubation (Fig. 7). Furthermore, holes and cracks appeared in some of the plastics incubated with bacteria (Fig. 6C, G-H). This might indicate that the communities are degrading the surface of the plastic at very slow rates, taking up the oxidation products at the same rate as they are set free from the LDPE. Thus, most changes in the surface structure were small or undetectable by our ATR-FTIR analysis (Fig. 5). It might also indicate that the communities are mainly utilizing compounds passively released by the plastics into the surrounding seawater, rather than actively degrading the polymer. Previously, it has been reported that the same LDPE sheets as used in this study release dissolved organic carbon (DOC) into the surrounding seawater, even when kept in the dark. This released DOC can subsequently be taken up by the marine microbial community ^58^. In our study it is not possible to determine whether the bacterial consortiums persisted by utilizing the LDPE polymer or the LDPE-associated additives, potentially leaching into the environment^59^. Recent evidence shows that certain bacteria can degrade specific plastic additives, some through pathways that also lead to fatty acid ß-oxidation ^60^. Utilization of additives, rather than the more stable LDPE polymeric structure, could also explain the decrease in weight of the LDPE pieces and cracking of the plastic surface, while also explaining the low signs of surface oxidation shown by the ATR-FTIR analysis. The degradation of these compounds in the marine environment is only poorly studied and to date there is no evidence that alkane-degraders like the ones enriched in our incubations can also degrade additives. However, due to the lack of knowledge of pathways involved in the biodegradation of most additives and the high variety of additive compounds, the possibility that certain organisms in our communities can utilize leaching LDPE-additives as energy and carbon source should not be disregarded.

There was also evidence of abiotic oxidation of plastics, as shown by low oxygen consumption in the controls (where microbes were absent) and in the ATR-FTIR spectra of the previously UV-weathered plastics. In our experiments abiotic oxidation of plastics was most obvious in the incubations with UV-weathered LDPE sheets (Fig. 5), where chemical compound classes were not detected in non-weathered LDPE and carbonyl groups decreased after six months of incubation even in the abiotic control ^61^. Usually, plastic biodegradation by microorganisms is more efficient when the polymer has previously undergone partial photo-oxidation ^62^. In our incubations this could not be detected, possibly due to the masking effect of the abiotic oxidation. This reinforces the fact that, if LDPE biodegradation occurs, it takes place at relatively slow rates, even when the plastic was previously UV-exposed.

Several heterotrophic bacteria like the organisms from Group R (Fig. 2) (*Rhodobactereaceae*) might have the ability to use alkanes as carbon and energy source. Whether they utilize plastics in the presence of other carbon sources, however, remains unknown. *Rhodobactereaceae* are typically among the most abundant families found in marine biofilms and are commonly enriched in plastic biofilms ^63–66^. Although the Group R MAGs were slightly enriched over time, they were already relatively abundant in the original community A0. They all harboured a gene encoding at least one alkane monooxygenase and the gene fadL, encoding a fatty acid transporter, was detected in all except one. In natural marine biofilms these organisms might have access to more easily accessible and metabolizable carbon and energy sources and therefore, might not participate in the uptake of LDPE-derived hydrocarbons. In contrast, obligatory hydrocarbonoclastic organisms, like *Alcanivorax borkumensis*, are specialized and dependent on degrading hydrocarbons ^31^. Group K organisms (Fig. 2) did exhibit several metabolic similarities and were closely related to *Alcanivorax borkumensis,* potentially occupying the same or a similar metabolic niche in the environment. For example, MAG 56, one of the most enriched MAGs, appears to lack proteins involved in the transport of sugars and amino acids into the cell. This might be an indication that, similar to *Alcanivorax borkumensis*, this bacterium relies exclusively on alkanes as a carbon source. However, we did not obtain completely closed genomes for any of the organisms in our incubations, which makes it impossible to determine with certainty whether they really lack these functions. Interestingly, even if present in relatively low abundances, the family *Alcanivoraceae* has been associated with plastics including PE in the marine environment ^4, 67, 68^. This might be an indication that these bacteria utilize PE as a carbon and energy source in the environment, consequently playing a role in the fate of plastics in the sea.

The lack of metagenomic data of the original communities B and C, does not allow to statistically determine MAGs and genes that were enriched after one year of incubation, when the major shifts in the community composition might have occurred. However, the development of communities A and C into relatively similar communities over time and the increase in abundance of taxa related to alkane and fatty acid degradation does support the hypothesis that alkane-degrading pathways are related to the degradation of PE-associated compounds. Taken together, we have identified a plethora of organisms, pathways and genes, other than the gene alkB, which might play an important role in the utilization of LDPE-associated compounds in the marine environment, thus providing a strong starting point for future studies addressing the mechanisms underlying microbial biodegradation of LDPE in the ocean.

## Materials and Methods

### Experimental setup

We collected three plastic pieces from the sea surface of the Northern Adriatic Sea, approximately 500 m off the coast of Rovinj, Croatia. Two were collected in November 2016 (plastics A and B) and one in August 2016 (plastic C). The plastics were immediately taken to the laboratory submerged in seawater (Fig. S6). A piece from each plastic was cut and frozen at −80°C for DNA analysis and another piece was taken for Raman spectroscopy to determine the plastic type. The three plastics were classified as polyethylene (PE). The remaining plastics were transferred to separate 1 L borosilicate bottles filled with artificial seawater (ASW) and amended with 2 µM of NaPO_4_ and 10 µM of NH_4_Cl (Table S9). The bottles were kept at room temperature (21°C-22.5°C)

After three weeks, the plastics were transferred to fresh 1 L bottles with ASW. This process was repeated seven times for plastic C and four times for plastics A and B. One small piece of each plastic type was then cut and incubated in a new bottle with ASW amended with inorganic nutrients (as described above) together with sterile low-density polyethylene (LDPE) pieces. The LDPE sheets were purchased from Goodfellow. Each bottle contained 10 LDPE squares (1×1 cm). After three weeks, one of these LDPE pieces was transferred to a new bottle with ASW, again together with new sterile LDPE pieces. This process was repeated for two years. After one and two years, a plastic sample from each of the incubations was taken and frozen at −80°C for metagenomics analyses. After two years, samples were also taken for respiration measurements.

All tools and bottles used were made of materials other than plastic and muffled or sterilized.

### DNA extraction and sequencing

DNA from plastics was extracted using a modified bead-beating approach combined with the Puregene Tissue DNA extraction kit (Qiagen, Valencia, CA) as previously described ^69^. Samples with sufficient amount of DNA were single-end sequenced (1 x 150bp) with Illumina NextSeq technology at Microsynth AG, Switzerland. These were plastic A right after collection (A0), after one year of incubation (A1), and after two years of incubation (A2), plastic C after one year and two years incubation (C1 and C2, respectively) and plastic B after one year of incubation (B1). All raw sequences files are available from the NCBI Sequence Read Archive (SRA) database (BioProject: PRJNA599974). Due to low DNA concentrations, samples from plastics B and C right after collection (samples B0 and C0) and sample B2 were not sequenced.

### Oxygen consumption measurements

At the end of the incubations after two years, one plastic piece from each of the incubations was transferred to a new bottle with ASW together with 16 sterile LDPE pieces. A second plastic piece from each of the incubations was incubated in a separate bottle with ASW with 16 glass pieces of 1 cm x 2.25 cm. As a control, 16 sterile LDPE pieces of dimensions 1.5 cm x 1.5 cm were incubated in a separate bottle with ASW (Control I). In order to let the biofilm develop on both glass and plastic without the influence of the old LDPE pieces, after one month, the plastics and glass were transferred to new ASW bottles and incubated for two additional months.

After two months, 15 pieces from each of the incubations were distributed among three BOD (biological oxygen demand) bottles of approximately 120 mL filled with ASW, resulting in triplicate bottles with five plastic pieces each. The same was done for the glass samples. One piece was also taken for bacterial abundance determination by adding it to 4% formaldehyde for 10 min and storing it at −80°C. Before transferring the glass and plastic to the BOD bottles for oxygen consumption measurements, one piece of glass and one of LDPE from each of the incubations were taken to determine cell abundance. The protocol for the oxygen and cell abundance measurements can be found in the Supplementary Information.

### Assessing the physical and chemical properties of the PE surface

To determine changes of the surface of PE sheets when incubated with the three communities after two years, we incubated one plastic colonized with one of the three communities in sterile ASW with three different types of PE pieces: (1) the same LDPE sheets as mentioned above; (2) the same LDPE sheets but previously UV-weathered (irradiated with UV radiation, wavelength 300-400 nm) under artificial conditions in a SUNSET CPS+ (Atlas-MTS) for two weeks, and (3) PE sheets from a pear packaging bag bought in a supermarket. Twenty pieces of each of the different types of PE sheets were incubated in 500 mL bottles, filled up to 300 mL with ASW and with one plastic already colonized with one of the three communities. A control bottle for each of the PE sheet types was prepared without the addition of an already colonized plastic. The bottles were kept at 21.5°C. After three months of incubation, one plastic sample from each bottle was collected for scanning electron microscopy (SEM) and at least quadruplicate samples were collected after six months for attenuated total reflection Fourier-transform infrared (ATR-FTIR) analysis. After eight months of incubation, at least five plastics were weighed from each incubation. Protocols for SEM and ATR-FTIR analysis can be found in the Supplementary Information.

### Statistical analysis and bioinformatics

ANOVA was performed to compare differences in total oxygen consumption and cell-specific oxygen consumption rates between samples with different communities and substrate. To assess differences between specific pairs of samples, post-hoc Tukey tests were performed. An ANOVA and post-hoc Tukey tests were also performed to compare the weight of plastics incubated with different communities after eight months of incubation. To identify genes enriched after two years of incubation in communities A and C, in comparison to the same communities incubated after one year, a differential expression analysis was performed using the DESeq2 analysis tools in R. Gene counts previously obtained with bbmap were used as input to the DESeq2 analysis. KEGG Annotated genes with significant differences in their fold-change in abundance, determined by the DESeq2 analysis (p-value < 0.05), between the two years were compared to the selected genes potentially involved in alkane and fatty acid biodegradation (Table S2). All statistical analyses and plots were done with the program R, unless otherwise stated.

Genomic assembly and annotation, read mapping, binning and further bioinformatics analysis can be found in the Supplementary Information.

### Genomic assembly and annotation

Read quality was assessed with fastqc. Overrepresented sequences were removed and reads were shortened to 149 bp in all samples using the adapterremoval program. After filtering and trimming, a total of 35,326,547, 15,541,923, 38,372,532, 3,180,678, 35,388,001 and 37,988,480 reads from samples A0, A1, A2, B1, C1 and C2, respectively, were co-assembled using the program Megahit ^70^ with default settings. Since we were aiming at identifying organisms and pathways enriched in the cultures, sample A0 was excluded from the co-assembly to decrease complexity.

Predicted genes were identified from the co-assembly using the Prodigal ^71^ software with default settings. Functional annotation of the predicted genes was performed with the online EggNOG-mapper v1 using Diamond as the sequence aligner with a maximum E-value cut-off of 10^-5^. A total of 2,500,984 predicted genes were recovered from the co-assembly from which 736,382 were annotated with the eggNOG emapper and from which 9,532 unique KO orthologies were identified. A total of 958 genes were enriched, i.e., increased in RPKM (reads per kilobase million) value over time (Table S5) of which 154 were classified as oxidoreductases (Table S6).

rRNA genes from each individual sample were assembled and further annotated through matam ^72^ using Silva_132_SSURef_Nr99 as the reference database. The identification of OTUs with matam is directly dependent on the correct assembly of one single gene (16S rRNA gene). Hence, it is likely to underestimate diversity in samples with a low number of reads by failing to assemble the gene of rare species. Metagenomes A1 and B1 had a significantly lower number of reads than the other samples. Therefore, to compare OTU diversity among all samples, the relative abundance and counts of OTUs were determined by the software mOTUs ^73^. This software classifies OTUs based on a group of 40 universal marker genes and hence, it is likely to be less biased towards identifying only relatively abundant OTUs than matam.

OTU diversity indexes Shannon and Simpson and OTU richness were calculated using the iNext R package ^74^. Coverage-based rarefaction was used to normalize all samples to a coverage of 84%, corresponding to the calculated coverage of the B1 metagenome which was the lowest of all samples.

### Read mapping

To obtain coverage information for assembled contigs and bins in all samples, the reads were mapped to the assembled contigs using the Burrows-Wheeler Aligner (BWA) ^75^. The mapped read counts, coverage and rpkm were extracted using bbmap/pileup.

### Binning

To recover individual genomes of the organisms enriched in the incubations, contigs from the co-assembly with a minimum of 1500 bp were binned using metabat ^76^ and maxbin^77^. In both cases, both tetra-nucleotide frequency and contig coverage in every individual sample were used for binning.

To obtain high-quality draft genomes, the resulting bins from each program were refined using metaWRAP ^78^. The resulting refined bins with completeness > 50% and contamination < 10% were used for further analysis. Hereafter, bins are called MAGs (metagenome assembled genomes).

Taxonomic classification of the MAGs was conducted via phylogenetic inference through marker genes using the software GTDB-Tk which uses the taxonomy as suggested by the Genome Taxonomy Database ^42^. We used Fasttree to build a phylogenetic tree with the multiple sequence alignment output by GTDB-Tk ^79^. MAG coverage in each sample was calculated by dividing the total number of bp of reads mapped to the MAG by the total length of all contigs and multiplying it by 10^6^ divided by the total number of bp of the sample. The count of mapped reads to the contigs of each MAG was obtained as described above. Gene calling of each MAG was predicted using Prodigal, sequences encoding alkB and P450 were retrieved by HMMER search (hmmscan) against a curated model downloaded from FunGene (http://fungene.cme.msu.edu/) with the threshold of domain coverage > 30%. The gene encoding CYP153 was further refined from the P450 gene with a customized hmm model with domain coverage > 75%. MAFFT ^80^ was used to build the alignment for both, the alkB and CYP153 gene. The phylogenetic tree was generated using RAxML^81^ under PROTGAMMALG model with 100 bootstraps. Tree visualization was carried using the Interactive Tree Of Life (iTOL) webtool ^82^.

### Enriched genes and metabolic pathways

Because oxidoreductase profiles have been found to be best suited to functionally characterize a microbial community ^83^, in addition to determining genes that increased in RPKM value over time, we specifically looked at the enriched oxidoreductases over time. Oxidoreductases were obtained by selecting KEGG orthology (KO) annotations obtained with EggNOG and classified as oxidoreductases (EC1). The EC numbers were obtained using KO to EC information provided by the ko2ec tool. The KO numbers of enriched genes were used as an input into the online KEGG Mapper reconstruction option and the complete enriched pathway modules were identified. We also analysed the relative abundance of selected genes (Table S2) encoding for enzymes previously suggested in the literature as being related to alkane degradation in all our metagenomes and also determined their presence in the individual MAGs.

## Data availability

All raw sequences files are available from the NCBI Sequence Read Archive (SRA) database (BioProject: PRJNA599974).

## Acknowledgments

We thank the Center for Marine Research of the Ruder Boskovic Institute, Croatia, for accommodating us during sampling, especially Dario, Mirjana, Ingrid, Paolo and Marino for all the support provided. We acknowledge Daniela Gruber for all the help provided during processing and analysis of the SEM samples. We also thank Miguel Guerreiro and Zihao Zhao for the assisting in bioinformatics issues. Financial support was provided by a Doctoral Fellowship (DOC scholarship) from the Austrian Academy of Sciences (ÖAW) awarded to MP, the ARTEMIS project of the Austrian Science Fund (P 28781-B21) to GJH and the research platform PLENTY financially supported by the University of Vienna, Austria.

## Author contributions

The study was designed by M.P. and G.J.H. Experiments were done by M.P. and data analysis was done by M.P. and Z.Z. ATR-FTIR analysis and interpretation were done by E.L. and K.K. The paper was written by M.P., Z.Z., K.K., E.L. and G.J.H.

## Competing interests

The authors declare no competing interests.

## References

1. Jambeck, J. R. et al. Plastic waste inputs from land into the ocean. Science. 347, 768– 771 (2015).

2. Amaral-Zettler, L. A., Zettler, E. R. & Mincer, T. J. Ecology of the plastisphere. Nat. Rev. Microbiol. 1–13 (2020) doi:10.1038/s41579-019-0308-0.

3. Zettler, E. R., Mincer, T. J. & Amaral-Zettler, L. Life in the ‘plastisphere’: Microbial communities on plastic marine debris. Environ. Sci. Technol. 47, 7137–7146 (2013).

4. Oberbeckmann, S., Osborn, A. M. & Duhaime, M. B. Microbes on a Bottle: Substrate, Season and Geography Influence Community Composition of Microbes Colonizing Marine Plastic Debris. PLoS One 11, e0159289 (2016).

5. Oberbeckmann, S., Löder, M. G. J. & Labrenz, M. Marine microplastic-associated biofilms–a review. Environ. Chem. 12, 551–562 (2015).

6. Amaral-Zettler, L. A. et al. The biogeography of the Plastisphere: implications for policy. Front. Ecol. Environ. 13, 541–546 (2015).

7. Oberbeckmann, S., Kreikemeyer, B. & Labrenz, M. Environmental Factors Support the Formation of Specific Bacterial Assemblages on Microplastics. Front. microbiol. 8, 2709 (2018).

8. Pinto, M., Langer, T. M., Hüffer, T., Hofmann, T. & Herndl, G. J. The composition of bacterial communities associated with plastic biofilms differs between different polymers and stages of biofilm succession. PLoS One 14, e0217165 (2019).

9. Kirstein, I. V et al. Dangerous hitchhikers? Evidence for potentially pathogenic Vibrio spp. on microplastic particles. Mar. Environ. Res. 120, 1–8 (2016).

10. Pinto, M. et al. Putative degraders of low-density polyethylene-derived compounds are ubiquitous members of plastic-associated bacterial communities in the marine environment. Environ. Microbiol. 22, 4779–4793 (2020).

11. Dussud, C. et al. Evidence of niche partitioning among bacteria living on plastics, organic particles and surrounding seawaters. Environ. Pollut. 236, 807–816 (2018).

12. Dussud, C. et al. Colonization of Non-biodegradable and Biodegradable Plastics by Marine Microorganisms. Front. Microbiol. 9, 1571 (2018).

13. Dudek, K. L., Cruz, B. N., Polidoro, B. & Neuer, S. Microbial colonization of microplastics in the Caribbean Sea. Limnol. Oceanogr. Lett. 5, 5–17 (2020).

14. Ogonowski, M. et al. Evidence for selective bacterial community structuring on microplastics. Environ. Microbiol. 20, 2796–2808 (2018).

15. Harrison, J. P., Schratzberger, M., Sapp, M. & Osborn, a. Rapid bacterial colonization of low-density polyethylene microplastics in coastal sediment microcosms. BMC Microbiol. 14, 232 (2014).

16. Oberbeckmann, S. et al. Genomic and proteomic profiles of biofilms on microplastics are decoupled from artificial surface properties. Environ. Microbiol. 23, 3099–3115 (2021).

17. Roager, L. & Sonnenschein, E. C. Bacterial Candidates for Colonization and Degradation of Marine Plastic Debris. Environ. Sci. Technol. 53, 11636–11643 (2019).

18. Syranidou, E. et al. Biodegradation of mixture of plastic films by tailored marine consortia. J. Hazard. Mater. 375, 33–42 (2019).

19. Geyer, R., Jambeck, J. R. & Law, K. L. Production, use, and fate of all plastics ever made. Sci. Adv. 3, e1700782 (2017).

20. Cózar, A. et al. Plastic debris in the open ocean. Proc. Natl. Acad. Sci. 111, 10239–10244 (2014).

21. Harshvardhan, K. & Jha, B. Biodegradation of low-density polyethylene by marine bacteria from pelagic waters, Arabian Sea, India. Mar. Pollut. Bull. 77, 100–106 (2013).

22. Sudhakar, M. et al. Biofouling and biodegradation of polyolefins in ocean waters. Polym. Degrad. Stab. 92, 1743–1752 (2007).

23. Devi, R. S., Ramya, R., Kannan, K., Antony, A. R. & Kannan, V. R. Investigation of biodegradation potentials of high density polyethylene degrading marine bacteria isolated from the coastal regions of Tamil Nadu, India. Mar. Pollut. Bull. 138, 549–560 (2019).

24. Kumari, A., Chaudhary, D. R. & Jha, B. Destabilization of polyethylene and polyvinylchloride structure by marine bacterial strain. Environ. Sci. Pollut. Res. 26, 1507–1516 (2019).

25. Muhonja, C. N., Makonde, H., Magoma, G. & Imbuga, M. Biodegradability of polyethylene by bacteria and fungi from Dandora dumpsite Nairobi-Kenya. PLoS One 13, e0198446 (2018).

26. Das, M. & Kumar, S. An approach to low-density polyethylene biodegradation by Bacillus amyloliquefaciens. 3 Biotech. 5, 81–86 (2014)

27. Yoon, M., Jeon, H. & Kim, M. Biodegradation of polyethylene by a soil bacterium and AlkB cloned recombinant cell. J Bioremed Biodegrad. 3, 1–8 (2012).

28. Syranidou, E. et al. Development of tailored indigenous marine consortia for the degradation of naturally weathered polyethylene films. PLoS One 12, e0183984 (2017).

29. Restrepo-Flórez, J.-M., Bassi, A. & Thompson, M. R. Microbial degradation and deterioration of polyethylene – A review. Int. Biodeterior. Biodegradation 88, 83–90 (2014).

30. Wang, W. & Shao, Z. Enzymes and genes involved in aerobic alkane degradation. Front. Microbiol. 4, 116 (2013).

31. Schneiker, S. et al. Genome sequence of the ubiquitous hydrocarbon-degrading marine bacterium Alcanivorax borkumensis. Nat. Biotechnol. 24, 997–1004 (2006).

32. Wang, W. & Shao, Z. The long-chain alkane metabolism network of Alcanivorax dieselolei. Nat. Commun. 5, 1–11 (2014).

33. Boonmak, C., Takahashi, Y. & Morikawa, M. Cloning and expression of three ladA-type alkane monooxygenase genes from an extremely thermophilic alkane-degrading bacterium Geobacillus thermoleovorans B23. Extremophiles 18, 515–523 (2014).

34. Allocati, N., Federici, L., Masulli, M. & Di Ilio, C. Glutathione transferases in bacteria. FEBS J. 276, 58–75 (2009).

35. Van Bogaert, I. N. A., Groeneboer, S., Saerens, K. & Soetaert, W. The role of cytochrome P450 monooxygenases in microbial fatty acid metabolism. FEBS J. 278, 206–221 (2011).

36. Gustafsson, M. C. U. et al. Expression, Purification, and Characterization of Bacillus subtilis Cytochromes P450 CYP102A2 and CYP102A3: Flavocytochrome Homologues of P450 BM3 from Bacillus megaterium. Biochemistry 43, 5474–5487 (2004).

37. Funhoff, E. G., Bauer, U., García-Rubio, I., Witholt, B. & van Beilen, J. B. CYP153A6, a Soluble P450 Oxygenase Catalyzing Terminal-Alkane Hydroxylation. J. Bacteriol. 188, 5220–5227 (2006).

38. Nie, Y. et al. Diverse alkane hydroxylase genes in microorganisms and environments. Sci. Rep. 4, 1–11 (2014).

39. Da Costa, J. P. et al. Degradation of polyethylene microplastics in seawater: Insights into the environmental degradation of polymers. J. Environ. Sci. Heal. Part A 53, 866– 875 (2018).

40. De Tender, C. et al. A review of microscopy and comparative molecular-based methods to characterize “Plastisphere” communities. Anal. Methods 9, 2132–2143 (2017).

41. Kim, S.-H. et al. Ketobacter alkanivorans gen. nov., sp. nov., an n-alkane-degrading bacterium isolated from seawater. Int. J. Syst. Evol. Microbiol. 68, 2258–2264 (2018).

42. Parks, D. H. et al. A standardized bacterial taxonomy based on genome phylogeny substantially revises the tree of life. Nat. Biotechnol. 36, 996–1004 (2018).

43. Kostka, J. E. et al. Hydrocarbon-Degrading Bacteria and the Bacterial Community Response in Gulf of Mexico Beach Sands Impacted by the Deepwater Horizon Oil Spill. Appl. Environ. Microbiol. 77, 7962–7974 (2011).

44. Kasai, Y. et al. Predominant growth of Alcanivorax strains in oil-contaminated and nutrient-supplemented sea water. Environ. Microbiol. 4, 141–147 (2002).

45. Schleheck, D. et al. Complete genome sequence of Parvibaculum lavamentivorans type strain (DS-1 T). Stand. Genomic Sci. 5, 298–310 (2011).

46. Lai, Q. et al. Parvibaculum indicum sp. nov., isolated from deep-sea water. Int. J. Syst. Evol. Microbiol. 61, 271–274 (2011).

47. Wang, L., Wang, W., Lai, Q. & Shao, Z. Gene diversity of CYP153A and AlkB alkane hydroxylases in oil-degrading bacteria isolated from the Atlantic Ocean. Environ. Microbiol. 12, 1230–1242 (2010).

48. Abbasian, F. et al. Microbial diversity and hydrocarbon degrading gene capacity of a crude oil field soil as determined by metagenomics analysis. Biotechnol. Prog. 32, 638–648 (2016).

49. Kummer, C., Schumann, P. & Stackebrandt, E. Gordonia alkanivorans sp. nov., isolated from tar-contaminated soil. Int. J. Syst. Evol. Microbiol. 49, 1513–1522 (1999).

50. Yakimov, M. M. et al. Alcanivorax borkumensis gen. nov., sp. nov., a new, hydrocarbon-degrading and surfactant-producing marine bacterium. Int. J. Syst. Evol. Microbiol. 48, 339–348 (1998).

51. Liu, C. & Shao, Z. Alcanivorax dieselolei sp. nov., a novel alkane-degrading bacterium isolated from sea water and deep-sea sediment. Int. J. Syst. Evol. Microbiol. 55, 1181– 1186 (2005).

52. Harwati, T. U., Kasai, Y., Kodama, Y., Susilaningsih, D. & Watanabe, K. Tranquillimonas alkanivorans gen. nov., sp. nov., an alkane-degrading bacterium isolated from Semarang Port in Indonesia. Int. J. Syst. Evol. Microbiol. 58, 2118–2121 (2008).

53. Nicdao, M. A. C. & Rivera, W. L. Two strains of Gordonia terrae isolated from used engine oil-contaminated soil utilize short-to long-chain n-alkanes. Philipp. Sci. Lett. 5, 1–10 (2012).

54. Gao, X. et al. Biodiversity and degradation potential of oil-degrading bacteria isolated from deep-sea sediments of South Mid-Atlantic Ridge. Mar. Pollut. Bull. 97, 373–380 (2015).

55. Guibert, L. M. et al. Diverse Bacterial Groups Contribute to the Alkane Degradation Potential of Chronically Polluted Subantarctic Coastal Sediments. Microb. Ecol. 71, 100–112 (2016).

56. Gerritse, J., Leslie, H. A., de Tender, C. A., Devriese, L. I. & Vethaak, A. D. Fragmentation of plastic objects in a laboratory seawater microcosm. Sci. Rep. 10, 1–16 (2020).

57. Sudhakar, M., Doble, M., Murthy, P. S. & Venkatesan, R. Marine microbe-mediated biodegradation of low- and high-density polyethylenes. Int. Biodeterior. Biodegradation 61, 203–213 (2008).

58. Romera-Castillo, C., Pinto, M., Langer, T. M., Álvarez-Salgado, X. A. & Herndl, G. J. Dissolved organic carbon leaching from plastics stimulates microbial activity in the ocean. Nat. Commun. 9, 1–7 (2018).

59. Suhrhoff, T. J. & Scholz-Böttcher, B. M. Qualitative impact of salinity, UV radiation and turbulence on leaching of organic plastic additives from four common plastics — A lab experiment. Mar. Pollut. Bull. 102, 84–94 (2016).

60. Wright, R. J., Bosch, R., Gibson, M. I. & Christie-Oleza, J. A. Plasticizer Degradation by Marine Bacterial Isolates: A Proteogenomic and Metabolomic Characterization. Environ. Sci. Technol. 54, 2244–2256 (2020).

61. Weiland, M., Daro, A. & David, C. Biodegradation of thermally oxidized polyethylene. Polym. Degrad. Stab. 48, 275–289 (1995).

62. Montazer, Z., Habibi-Najafi, M. B., Mohebbi, M. & Oromiehei, A. Microbial Degradation of UV-Pretreated Low-Density Polyethylene Films by Novel Polyethylene-Degrading Bacteria Isolated from Plastic-Dump Soil. J. Polym. Environ. 26, 3613–3625 (2018).

63. Oberbeckmann, S. & Labrenz, M. Marine Microbial Assemblages on Microplastics: Diversity, Adaptation, and Role in Degradation. Ann. Rev. Mar. Sci. (2019) doi:10.1146/annurev-marine-010419-010633.

64. Kesy, K., Oberbeckmann, S., Kreikemeyer, B. & Labrenz, M. Spatial Environmental Heterogeneity Determines Young Biofilm Assemblages on Microplastics in Baltic Sea Mesocosms. Front. Microbiol. 10, 1665 (2019).

65. Bryant, J. A. et al. Diversity and Activity of Communities Inhabiting Plastic Debris in the North Pacific Gyre. mSystems 1, e00024–16 (2016).

66. Debroas, D., Mone, A. & Ter Halle, A. Plastics in the North Atlantic garbage patch: A boat-microbe for hitchhikers and plastic degraders. Sci. Total Environ. 599–600, 1222–1232 (2017).

67. Oberbeckmann, S., Loeder, M. G. J., Gerdts, G. & Osborn, A. M. Spatial and seasonal variation in diversity and structure of microbial biofilms on marine plastics in Northern European waters. FEMS Microbiol. Ecol. 90, 478–492 (2014).

68. Delacuvellerie, A., Cyriaque, V., Gobert, S., Benali, S. & Wattiez, R. The plastisphere in marine ecosystem hosts potential specific microbial degraders including Alcanivorax borkumensis as a key player for the low-density polyethylene degradation. J. Hazard. Mater. 380, 120899 (2019).

69. Debeljak, P. et al. Extracting DNA from ocean microplastics: a method comparison study. Anal. Methods 9, 1521–1526 (2017).

70. Li, D., Liu, C.-M., Luo, R., Sadakane, K. & Lam, T.-W. MEGAHIT: an ultra-fast single-node solution for large and complex metagenomics assembly via succinct de Bruijn graph. Bioinformatics 31, 1674–1676 (2015).

71. Hyatt, D. et al. Prodigal: prokaryotic gene recognition and translation initiation site identification. BMC Bioinformatics 11, 1–11 (2010).

72. Pericard, P., Dufresne, Y., Couderc, L., Blanquart, S. & Touzet, H. MATAM: reconstruction of phylogenetic marker genes from short sequencing reads in metagenomes. Bioinformatics 34, 585–591 (2017).

73. Milanese, A. et al. Microbial abundance, activity and population genomic profiling with mOTUs2. Nat. Commun. 10, 1–11 (2019).

74. Hsieh, T. C., Ma, K. H. & Chao, A. iNEXT : an R package for rarefaction and extrapolation of species diversity ( Hill numbers). 1451–1456 (2016) doi:10.1111/2041-210X.12613.

75. Li, H. & Durbin, R. Fast and accurate short read alignment with Burrows–Wheeler transform. bioinformatics 25, 1754–1760 (2009).

76. Kang, D. D., Froula, J., Egan, R. & Wang, Z. MetaBAT, an efficient tool for accurately reconstructing single genomes from complex microbial communities. PeerJ 3, e1165– e1165 (2015).

77. Wu, Y.-W., Tang, Y.-H., Tringe, S. G., Simmons, B. A. & Singer, S. W. MaxBin: an automated binning method to recover individual genomes from metagenomes using an expectation-maximization algorithm. Microbiome 2, 1–18 (2014).

78. Uritskiy, G. V, DiRuggiero, J. & Taylor, J. MetaWRAP—a flexible pipeline for genome-resolved metagenomic data analysis. Microbiome 6, 1–13 (2018).

79. Chaumeil, P.-A., Mussig, A. J., Hugenholtz, P. & Parks, D. H. GTDB-Tk: a toolkit to classify genomes with the Genome Taxonomy Database. Bioinformatics, 1925–1927 (2019)

80. Katoh, K. & Standley, D. M. MAFFT multiple sequence alignment software version 7: improvements in performance and usability. Mol. Biol. Evol. 30, 772–780 (2013).

81. Stamatakis, A. RAxML version 8: a tool for phylogenetic analysis and post-analysis of large phylogenies. Bioinformatics 30, 1312–1313 (2014).

82. Letunic, I. & Bork, P. Interactive tree of life (iTOL) v3: an online tool for the display and annotation of phylogenetic and other trees. Nucleic Acids Res. 44, 242–245 (2016).

83. Ramírez-Flandes, S., González, B. & Ulloa, O. Redox traits characterize the organization of global microbial communities. Proc. Natl. Acad. Sci. 116, 3630–3635 (2019).

